# Cell-type-specific Rab–Munc13-4 coupling defines divergent exosome fusion mechanisms and post-docking fusion pore opening

**DOI:** 10.64898/2026.11.24.769064

**Authors:** Sang Su Woo

## Abstract

Munc13-4 operates as an intracellular Ca^2+^ sensor for regulated secretion in both immune and cancer cells. Its capacity to physically engage multiple Rab GTPases and SNARE proteins places it at the center of membrane reorganization upon intracellular Ca^2+^ elevation. Here, we define the mode of action of Munc13-4 during Ca^2+^-evoked exosome secretion in RBL-2H3 mast cells. We show that Munc13-4 coupling to Rab27a is essential to retain Munc13-4 on secretory membranes for exosome release in mast cells, contrasting with cancer cells where the Munc13-4–Rab11 axis predominates. Using simultaneous epifluorescence and total internal reflection fluorescence (TIRF) microscopy validated by high-pressure freezing electron microscopy, fluorescence recovery after photobleaching (FRAP), and staged acute inhibition with Bexin-1, we demonstrate that Munc13-4 acts at the terminal post-docking stage of multivesicular body (MVB) exocytosis: acute Munc13-4 inhibition after vacuole formation leaves PM contact intact but abolishes fusion pore opening and destabilizes nascent pores. Moreover, depleting Rab27a while elevating Rab11 redirects Munc13-4 onto Rab11^+^ compartments in mast cells, recapitulating the cancer-cell phenotype. Together, our findings support a rapid-exchange, abundance-weighted Rab partitioning model that tempers cellular Ca^2+^ responsiveness and governs cell-type-specific exosome fusion pathways, providing a mechanistic rationale for context-selective modulation of pathological exosome release.

## Introduction

Whereas the biogenesis of intraluminal vesicles (ILVs) within multivesicular bodies (MVBs) has been extensively characterized (Trajkovic et al., 2008; Cocucci and Meldolesi, 2015; Lo Cicero et al., 2015; Jackson et al., 2017; Stahl and Raposo, 2018; van Niel et al., 2018), the molecular mechanisms governing terminal MVB exocytosis at the plasma membrane (PM) remain incompletely defined. Previous studies have implicated distinct, cell-type-specific Rab GTPases in MVB tethering and docking at the PM: Rab27a and Rab27b regulate MVB exocytosis by recruiting their effectors Slp4 and Slac2b and remodeling cortical actin in HeLa cells (Ostrowski et al., 2010); Rab11 and Rab35 mediate MVB docking in erythroleukemia, reticulocyte, and oligodendroglial cells (Savina et al., 2005; Hsu et al., 2010); and the small GTPase RAL-1 controls MVB biogenesis and PM docking in *Caenorhabditis elegans* (Hyenne et al., 2015). In activated immune cells, intracellular Ca^2+^ elevation triggers the formation of large endolysosomal vacuoles linked to exosome secretion (Savina et al., 2003; Omari et al., 2024). However, continuous real-time visualization spanning vacuole assembly, PM docking, fusion pore opening, and exosome expulsion remains rare, and why MVBs exploit divergent Rab GTPase modules across cell types is poorly understood.

Mast cells actively secrete exosomes carrying functional mRNAs, miRNAs, and immunoregulatory proteins upon cytoplasmic Ca^2+^ elevation, contributing critically to host immunity (Valadi et al., 2007; Vukman et al., 2017). While vesicle priming—defined here as the Ca^2+^-dependent maturation of pre-docked vesicles into a fusion-competent state—is well characterized in neurons via Munc13-1/2, Complexin, and CAPS, the priming machinery in immune cells is less defined. RBL-2H3 mast cells lack canonical Syntaxin-1 and instead rely on Syntaxin-11 stabilized by its chaperone Munc18-2, an axis essential for plasma membrane localization and terminal granule fusion in cytotoxic lymphocytes (Dieckmann et al., 2015). Given low Complexin II abundance and undefined CAPS regulators in mast cells, the C2-domain Ca^2+^ sensor Munc13-4—whose loss-of-function mutations cause familial hemophagocytic lymphohistiocytosis type 3 (FHL3; Feldmann et al., 2003)—likely serves as a primary Ca^2+^-dependent priming factor. Munc13-4 interacts with Rab and SNARE proteins to promote secretory granule (SG) tethering, endosomal remodeling, and Ca^2+^-stimulated liposome fusion (Neeft et al., 2005; Elstak et al., 2011; Boswell et al., 2012; Chicka et al., 2016; He et al., 2016; Johnson et al., 2016; Woo et al., 2017). Previously, we showed that Munc13-4 acts as a Ca^2+^ sensor driving homotypic Rab7^+^ SG fusion to generate enlarged exocytic MVBs (vacuoles) in RBL-2H3 cells (Woo et al., 2017), whereas in cancer cells Munc13-4 operates as a Rab11 effector mediating Ca^2+^-stimulated CD63^+^ exosome release (Messenger et al., 2018).

Despite these advances, whether Munc13-4 is directly required at the terminal post-docking stage of MVB–PM fusion pore opening in live cells has remained unresolved. Conventional supernatant assays measure only accumulated exosome end-products, while pH-sensitive reporters such as CD63-pHluorin detect terminal luminal neutralization without resolving upstream vacuole dynamics, effector retention at the fusion site, or stalled post-docking intermediates (Messenger et al., 2018; Verweij et al., 2018; Lu et al., 2018). Furthermore, although acute chemical inhibition of Munc13-4 by Bexin-1 blocks late-stage SG exocytosis (Bruinsma et al., 2018), its impact on pre-formed, PM-apposed MVBs—and how intracellular Rab stoichiometry governs Munc13-4 targeting across cell types—has not been directly tested.

Here, we establish a real-time imaging framework resolving vacuole assembly, convergent cargo incorporation, PM docking, and micron-scale fusion pore opening in RBL-2H3 mast cells using simultaneous epifluorescence and TIRF microscopy corroborated by high-pressure freezing electron microscopy (HPF-EM). Combining knockdown approaches with staged acute Bexin-1 application, we demonstrate that Munc13-4 is specifically required after vacuole formation and PM contact to drive fusion pore opening and stabilization. Finally, single-cell FRAP kinetics, *in situ* cross-linking immunoprecipitation, and Rab27a-depletion/Rab11-elevation experiments establish an abundance-weighted Rab partitioning model that explains both cell-type-specific routing of Munc13-4 and the graded Ca^2+^ sensitivity of intact secretory cells.

## Results

### Direct live-cell visualization of Ca²⁺-evoked vacuole–PM fusion pore opening and CD63⁺ exosome release

To visualize vacuoles and exosomes simultaneously in live RBL-2H3 cells, C-terminally tagged CD63 (CD63-mKate2) was co-expressed with EGFP-Munc13-4, and events in the cell body (epifluorescence, Epi) and basal PM (TIRF) were recorded concurrently every 20 s at 37°C before and after 2 μM ionomycin stimulation (**Fig. 1A**). In resting cells, CD63-mKate2 appeared as small rings or puncta that partially overlapped with EGFP-Munc13-4 in the TIRF field (**Fig. 1A**, 0 min), representing pre-docked SGs. Following ionomycin addition, pre-docked SGs rapidly underwent exocytosis while cytoplasmic vesicles in perinuclear regions merged homotypically to form large vacuoles (>2.5 μm diameter), accompanied by accumulation of CD63-mKate2 within the vacuole lumen (**Fig. 1A**, 12 min Epi inset; Woo et al., 2017).

**Figure 1.**
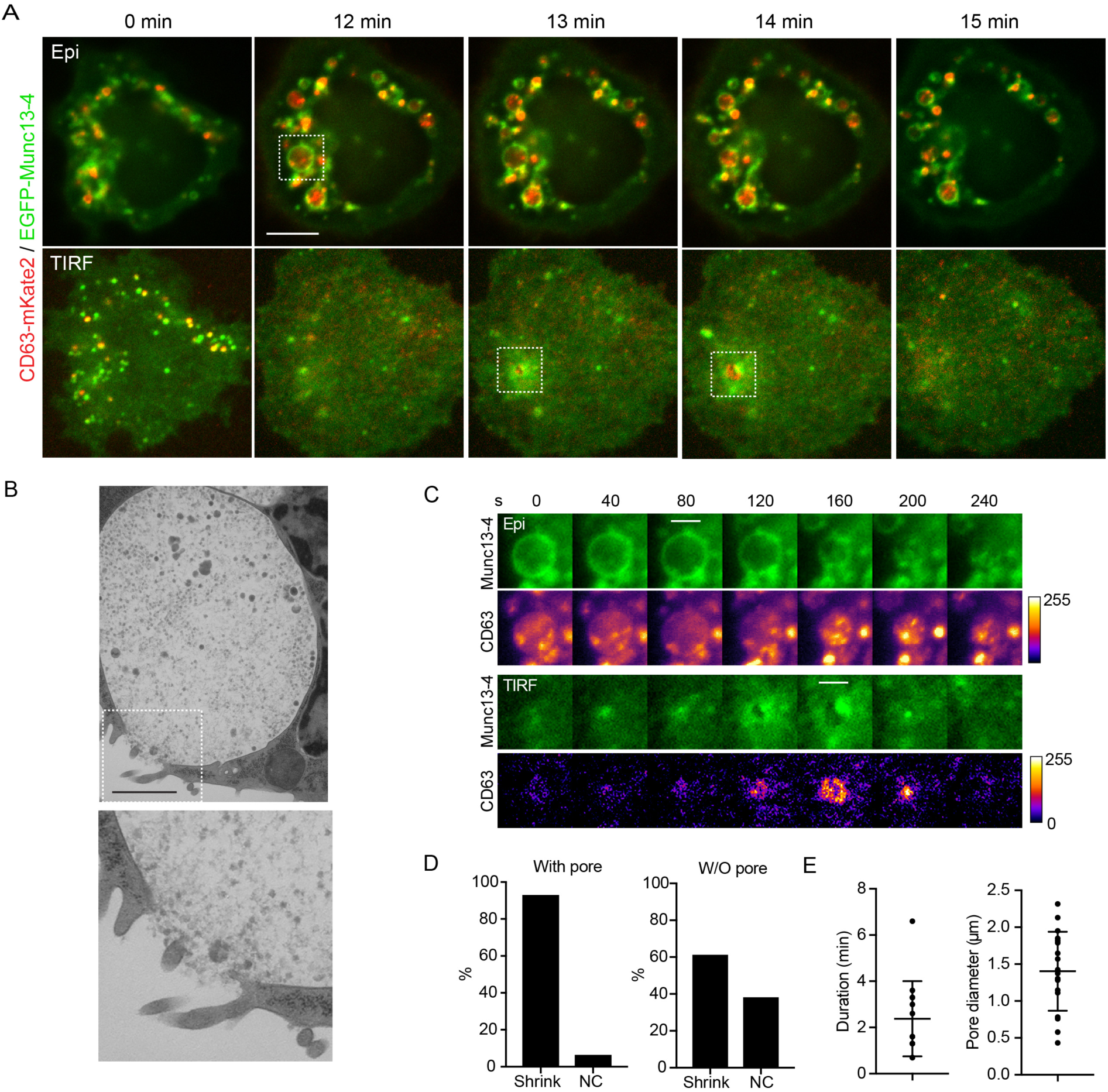
Direct imaging of CD63^+^ exosome release from vacuoles. **A.** Time-lapse images of vacuole exocytosis captured by simultaneous epi-TIRF imaging. RBL-2H3 cells co-expressing CD63-mKate2 (red) and EGFP-Munc13-4 (green) were treated with 2 μM ionomycin and imaged every 20 s. Scale bar, 10 μm. **B.** High-pressure freezing/freeze-substitution electron micrographs showing exosome release through a vacuole–PM fusion opening (fusion pore). Bottom image is a magnified inset of the boxed region. Scale bar, 1 μm. **C.** Time-lapse montages (40-s intervals) for the inset in A. CD63-mKate2 was pseudo-colored with a fire heatmap LUT (right-side scale bars, 0–255). Scale bars: Epi, 3 μm; TIRF, 1 μm. **D.** Fraction of vacuoles undergoing shrinkage (’Shrink’) versus no change in size (’NC’). Among 28 vacuoles analyzed by visual inspection, 15 showed a basal TIRF fusion pore (’With pore’; 93% shrank upon PM fusion), whereas 13 did not (’W/O pore’; 62% shrank). **E.** Vacuole fusion pore duration (min) and peak pore diameter (μm). Lines denote mean ± SEM (*n* = 14 fusion pores).

Migrating vacuoles eventually anchored at discrete PM sites and fused with the PM (**Fig. 1A**, 13–14 min TIRF insets). Upon PM contact, EGFP-Munc13-4 fluorescence intensified at the TIRF footprint and subsequently spread radially along the PM, leaving a central Munc13-4-depleted zone filled with luminal CD63-mKate2 puncta (**Fig. 1A, C**). Because this ring-like geometry coincided with luminal content expulsion, it reported the opening and lateral dilation of an MVB–PM fusion pore (Woo et al., 2017). Time-lapse montages at 40-s intervals directly resolved outward diffusion of luminal CD63-mKate2 puncta through the expanding pore (**Fig. 1C**).

The presence of micron-scale MVB–PM fusion openings and the release of 50–120 nm intraluminal vesicles were independently confirmed by transmission electron microscopy of samples prepared by high-pressure freezing and freeze substitution (HPF-EM; **Fig. 1B**). Luminal CD63-mKate2 puncta exhibited a full width at half maximum (FWHM) of 893 ± 139 nm (mean ± SD, *n* = 3), matching the dimensions of aggregated exosomes observed by HPF-EM (**Supplementary Fig. S1A**). Released extracellular vesicles comprised two morphological subpopulations in EM micrographs: a homogeneous smaller group (∼70 nm diameter) and a more variable larger group (∼100 nm diameter). To our knowledge, these observations provide the first direct live-cell fluorescence visualization of CD63^+^ exosome expulsion through a resolved MVB–PM fusion pore.

Quantitative tracking of individual vacuoles (*n* = 28 vacuoles) showed that the majority underwent exocytosis (**Fig. 1D, E**). Specifically, 15 of 28 vacuoles (54%) displayed a directly visible fusion pore in the basal TIRF plane (’With pore’), among which 93% (14/15) shrank immediately upon fusion (**Fig. 1D**, left), establishing vacuole shrinkage as a reliable morphological readout of exocytosis. Among the remaining 13 vacuoles (46%) without a visible basal pore (’W/O pore’), 62% (8/13; corresponding to an additional 29% of total vacuoles) nevertheless shrank acutely (**Fig. 1D**, right), consistent with fusion pore opening at apical or lateral PM regions outside the TIRF field. Overall, 79% (22/28) of vacuoles underwent exocytic shrinkage, and ∼54–82% are estimated to undergo exocytosis. Resolved fusion pores (*n* = 14) expanded to peak diameters of 1.4 ± 0.5 μm and persisted for 2.38 ± 1.63 min before closing or dissipating (**Fig. 1E**).

### Maturing vacuoles dynamically incorporate distinct exosome cargoes via convergent trafficking routes

Live-cell tracking also captured continuous homotypic fusion and fission cycles during vacuole maturation prior to PM fusion (**Fig. 2A**; comparable to morphological remodeling noted in Omari et al., 2024 and Woo et al., 2017). Across 14 examined cells, 57% formed at least one vacuole (*n* = 28 vacuoles from 8 cells across 4 experiments), exhibiting an average vacuole lifespan of 14.27 ± 0.93 min (**Fig. 2A**, inset). Initial homotypic fusion occurred with a latency of 6.46 ± 0.93 min after ionomycin addition, whereas vacuole exocytosis occurred at 12.31 ± 1.11 min. In resting cells, ∼60–67% of ring-shaped CD63 structures colocalized with Munc13-4 on their limiting membrane (**Fig. 2B**, arrow; **Supplementary Fig. S2A**), while endosomal CD63 overlapped extensively with Rab7^+^ (∼73%) and Rab11^+^ (∼48%) compartments (**Fig. 2B**, arrowheads; **Supplementary Fig. S2B**). Upon ionomycin stimulation, early SG exocytosis transiently inserted CD63 into the basal PM, increasing PM CD63 intensity by ∼40% within 5 min (**Supplementary Fig. S1B**), whereas intracellular CD63 vesicles coalesced into enlarging vacuoles (**Fig. 2C**). Radial intensity profiling and time-lapse imaging demonstrated progressive luminal accumulation and *de novo* formation of CD63^+^ puncta inside growing vacuoles (**Fig. 2C, D**), indicating active ILV generation during vacuole maturation (**Fig. 2E**).

**Figure 2.**
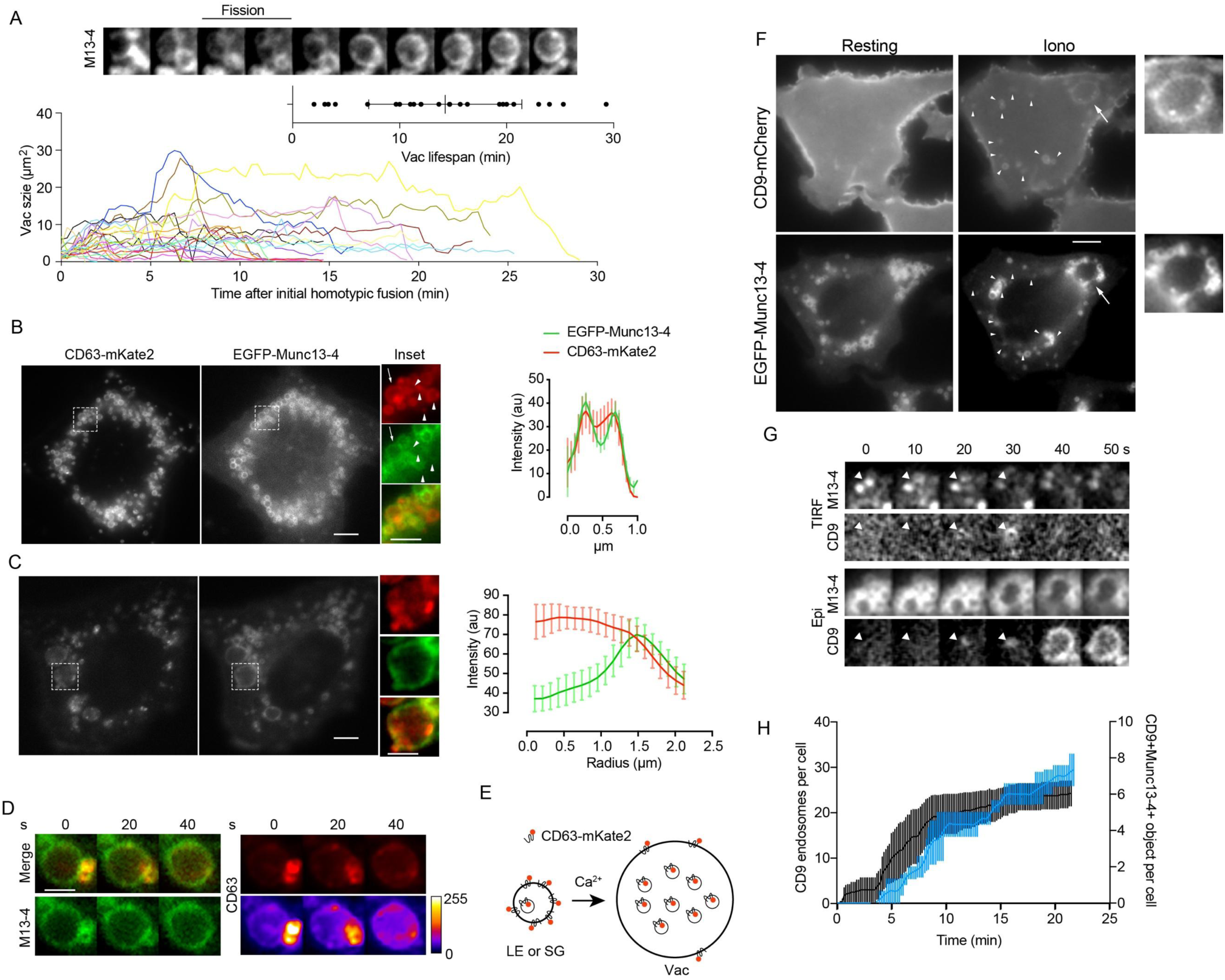
Vacuole maturation dynamically incorporates distinct exosome cargoes. **A.** Top: Time-lapse montage showing vacuole fission and re-fusion marked by EGFP-Munc13-4. Bottom: Growth dynamics of individual color-coded vacuoles plotted over time after initial homotypic fusion. Inset dot plot denotes vacuole lifespan (mean ± SEM; *n* = 28 vacuoles from 8 cells across 4 independent experiments; 57% of 14 examined cells formed at least one vacuole, a higher frequency than fixed-cell ICC results [28%], Woo et al., 2017). **B.** CD63-mKate2 in SGs and endosomes in resting RBL-2H3 cells. Arrow, Munc13-4^+^/ CD63^+^ SG; arrowheads, CD63^+^ endosomes. Right: Line-scan intensity profile showing CD63 overlap with Munc13-4 on the limiting membrane (mean ± SEM; *n* = 18 SGs from 3 cells). Scale bar, 5 μm. **C.** CD63-mKate2 accumulation in the vacuole lumen after ionomycin stimulation. Right: Radial intensity profile across vacuoles (mean ± SEM; *n* = 6 vacuoles from 3 cells); the EGFP-Munc13-4 peak marks the limiting membrane. Scale bar, 5 μm (inset, 3 μm). **D.** *De novo* appearance of luminal CD63^+^ ILV puncta in enlarging vacuoles (20-s intervals; bottom CD63 panels pseudo-colored with a heatmap LUT). Scale bar, 3 μm. **E.** Schematic representation of CD63 localization in SGs/late endosomes (LE) versus maturing vacuoles (Vac). **F.** Live-cell snapshots of cells co-expressing CD9-mCherry and EGFP-Munc13-4 under resting and ionomycin-treated conditions. Arrow, vacuole (enlarged at right); arrowheads, CD9^+^ endosomes. Scale bar, 5 μm. **G.** Dual TIRF/Epi montage (10-s intervals) showing PM-derived CD9^+^ endosome formation following local SG exocytosis (arrowheads) and subsequent delivery to vacuoles. **H.** Kinetics of CD9^+^ endosome formation (black, left axis) and emergence of CD9^+^/Munc13-4^+^ objects per cell (blue, right axis). Data indicate mean ± SEM (*n* = 3 cells from 2 experiments).

We next tested whether enlarging vacuoles incorporate distinct exosome cargoes originating from separate cellular membranes. Although CD9 is a biochemically established exosome marker, its trafficking route in mast cells was unknown (Köberle et al., 2012). Co-expression of CD9-mCherry with EGFP-Munc13-4 revealed that, in contrast to endogranular CD63, CD9 resided almost exclusively at the PM in resting cells and translocated onto vacuoles after ionomycin stimulation, appearing both on the Munc13-4^+^ limiting membrane and as luminal puncta (**Fig. 2F**). Dual epi-TIRF time-lapse sequences showed that small CD9^+^ endocytic vesicles emerged from the PM immediately following local SG exocytosis and subsequently merged with nascent Munc13-4^+^ vacuoles (**Fig. 2G**), explaining the delayed rise of CD9^+^/Munc13-4^+^ vacuoles relative to initial CD9 endosome formation (**Fig. 2H**). Time-course secretion assays confirmed Ca^2+^-dependent release of CD9-GFP (∼5–7% within 15–20 min, with an ∼8 min onset latency matching vacuole exocytosis) while cytosolic GFP control was not released (**Supplementary Fig. S2C**). Because no direct PM shedding of CD9 vesicles was observed at ∼100 nm optical resolution, these data establish that PM-endocytosed CD9 and endogranular CD63 converge onto maturing vacuoles prior to exosome secretion.

### Munc13-4 is required at a post-docking step for vacuole–PM fusion pore opening and stabilization

To test the requirement for Munc13-4 in mast cell exosome secretion, we depleted Munc13-4 by lentiviral shRNA transduction (**Supplementary Fig. S3A**) and quantified Ca^2+^-stimulated release using complementary assays. In 96-well fluorometric assays, Munc13-4 knockdown reduced ionomycin-evoked secretion of both CD9-EGFP and CD63-mKate2 by ∼80% without affecting basal release (**Fig. 3A**). Consistently, slot-blot immunodetection of differential-sedimentation supernatants showed robust Ca^2+^-triggered secretion of endogenous CD63 and CD9—devoid of *cis*-Golgi GM130 contamination—which was strongly abolished by Munc13-4 knockdown (**Fig. 3B**). Equilibrium sucrose density gradient fractionation (20%/40% interface) further verified that secreted CD63 and CD9 resided on canonical exosomal vesicles whose stimulated release depended on Munc13-4 (**Supplementary Fig. S3B**).

**Figure 3.**
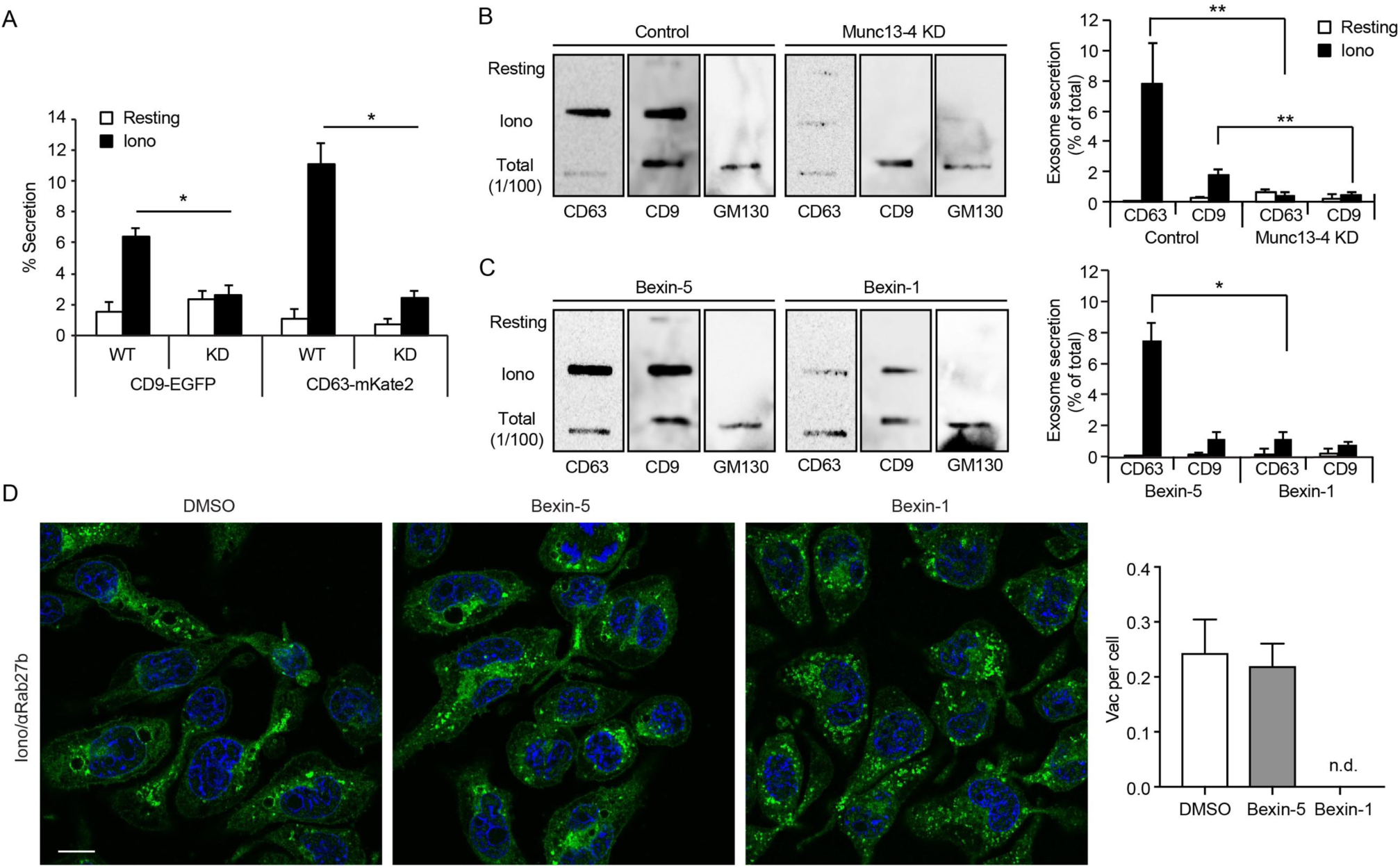
Munc13-4 is required for Ca^2+^-stimulated exosome secretion in RBL-2H3 cells. **A.** Effect of Munc13-4 knockdown (KD) on Ca^2+^-stimulated secretion of CD9-EGFP and CD63-mKate2 in resting or ionomycin-treated cells determined by 96-well fluorometric assay. Released CD9 and CD63 were calculated as % of total cellular fluorescence (Woo et al., 2017). Bars denote mean ± SEM (*n* = 3; *, *P* < 0.05). **B.** Complementary slot-blot assay of endogenous exosome secretion from control (WT) or Munc13-4 KD cells treated with 2.5 μM ionomycin for 20 min at 37°C (Messenger et al., 2018). The *cis*-Golgi marker GM130 was probed as a negative control for intracellular membrane contamination alongside 1/100 total cell lysates. Right: Densitometric quantification (% of total; mean ± SEM, *n* = 3; **, *P* < 0.01). **C.** Effect of the small-molecule Munc13-4 inhibitor Bexin-1 (10 μM) versus its inactive sham control Bexin-5 (10 μM), applied 10 min prior to ionomycin stimulation, measured as in B. Right: Quantification (mean ± SEM, *n* = 4; *, *P* < 0.05). **D.** Bexin-1 inhibits Ca^2+^-evoked vacuole formation. Cells pre-treated for 10 min with DMSO, Bexin-5, or Bexin-1 prior to ionomycin stimulation were immunostained for Rab27b (green) and Hoechst 33342 (blue) to quantify enlarged vacuoles (>2.5 μm diameter). Scale bar, 10 μm. Right: Vacuole frequency per cell (mean ± SEM; *n* = 94, 145, and 75 cells for DMSO, Bexin-5, and Bexin-1, respectively; n.d., not detected).

We next employed Bexin-1, an acute small-molecule inhibitor of Munc13-4-mediated membrane fusion, paired with its inactive structural analog Bexin-5 as a negative control (Bruinsma et al., 2018). Pre-incubating RBL-2H3 cells for 10 min with suboptimal Bexin-1 (10 μM), but not Bexin-5, suppressed ionomycin-stimulated endogenous CD63 and CD9 exosome secretion to levels comparable to Munc13-4 knockdown without perturbing basal secretion or causing GM130 leakage (**Fig. 3C**). Likewise, 10-min pre-treatment with Bexin-1 completely blocked the formation of enlarged Rab27b^+^ vacuoles (>2.5 μm diameter), whereas Bexin-5 had no effect relative to DMSO vehicle (**Fig. 3D**).

Because pre-treatment halts upstream homotypic SG fusion, we exploited the rapid onset of Bexin-1 action (2–3 min; Bruinsma et al., 2018) in a staged live-cell protocol to isolate the terminal vacuole–PM fusion step (**Fig. 4A**). Cells expressing EGFP-Munc13-4 were first stimulated with 2 μM ionomycin until initial homotypic vacuole formation was visually confirmed, at which point 10 μM Bexin-5 or Bexin-1 was acutely injected while monitoring Epi and TIRF channels (**Fig. 4A, B**). Vacuoles exposed to control Bexin-5 continued enlarging and opened resolved TIRF fusion pores within 4.4 ± 2.3 min after compound addition (*n* = 7 vacuoles; **Fig. 4B, C**). By contrast, acute Bexin-1 addition completely abolished fusion pore opening and lateral Munc13-4 dispersion across the PM (0 of 10 vacuoles; n.d.), even though pre-formed vacuoles remained stably apposed to the basal TIRF footprint (**Fig. 4B, C**). Furthermore, quantitative analysis of resolved pore-opening events across experiments revealed that even when pores occasionally initiated in the presence of Bexin-1, their open lifetimes were sharply curtailed (median duration 20 s, IQR 20–55 s, *n* = 4 events) compared with control pores (median 160 s, IQR 80–260 s, *n* = 10 events; **Fig. 4D**). Accordingly, the fraction of ultra-short, prematurely collapsing pores (≤20 s) rose from 10% (1/10) in control events to 75% (3/4) upon Bexin-1 treatment (two-sided Fisher’s exact test, *P* = 0.041; **Fig. 4E**), indicating that Munc13-4 is required both to trigger pore opening and to stabilize the dilating fusion pore.

**Figure 4.**
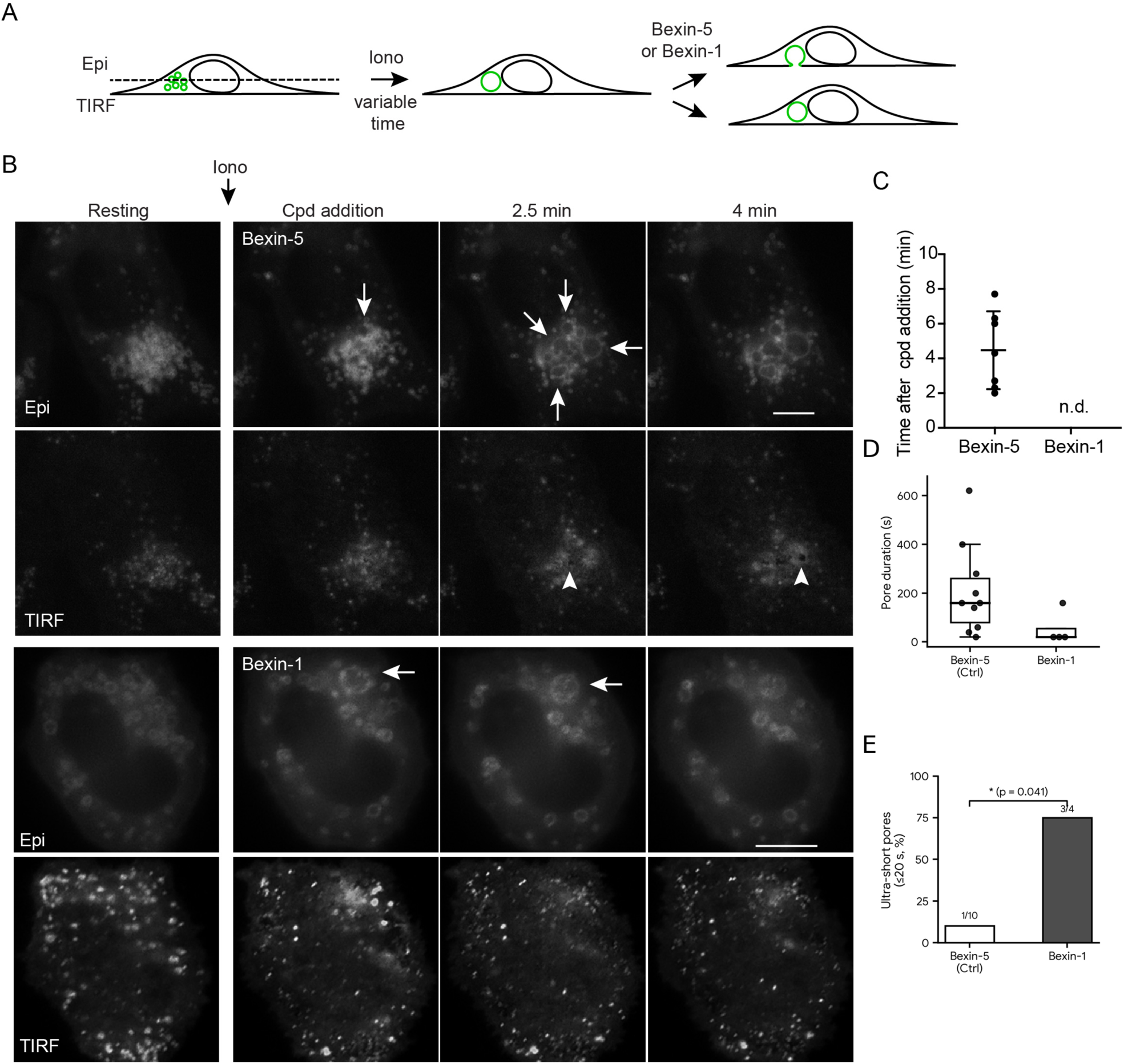
Munc13-4 is required for vacuole–PM fusion pore opening and stabilization. **A.** Experimental schematic for testing acute Bexin-1 inhibition on vacuole–PM fusion. RBL-2H3 cells expressing EGFP-Munc13-4 (green) were stimulated with 2 μM ionomycin at 37°C. As soon as initial homotypic secretory granule (SG) fusion was observed, 10 μM Bexin-1 or its inactive analog Bexin-5 (sham control) was acutely added, and dual-plane time-lapse images of the cell body (Epi) and basal plasma membrane (TIRF) were continuously acquired. **B.** Sequential addition of Bexin-1 inhibits vacuole–PM fusion. Representative Epi and TIRF time-lapse images show vacuole formation and exocytosis marked by EGFP-Munc13-4 fluorescence. White arrows indicate vacuoles in Epi panels; arrowheads in Bexin-5 TIRF panels indicate a vacuole–PM fusion event with a dilating fusion pore. The vacuole in the Bexin-1-treated cell remains apposed to the PM without opening a fusion pore. Note that the Bexin-5-treated vacuole continues to enlarge, whereas the Bexin-1-treated vacuole does not. Scale bars, 10 μm. **C.** Time from compound addition to fusion pore opening in the staged-addition assay shown in B. Lines indicate mean ± SD (Bexin-5, *n* = 7 vacuoles from 3 cells, 2 experiments; Bexin-1, *n* = 10 vacuoles from 6 cells, 4 experiments; n.d., not detected). **D.** Distribution of fusion pore open durations (s) for resolved pore-opening events in Bexin-5 (Ctrl) (median 160 s, IQR 80–260 s; *n* = 10 events) versus Bexin-1-treated cells (median 20 s, IQR 20–55 s; *n* = 4 events). Boxes represent the interquartile range (IQR) with the horizontal line denoting the median, and individual events are overlaid as dots. **E.** Fraction of ultra-short, prematurely collapsing fusion pores (≤20 s) among resolved pore-opening events. Bexin-1 markedly increased the proportion of ultra-short pores from 10% (1/10) in Bexin-5 (Ctrl) to 75% (3/4) (*, *P* = 0.041, two-sided Fisher’s exact test).

### Rab27a retains Munc13-4 on mast cell membranes, whereas Rab27a loss enables cancer-like switching to Rab11

Because Munc13-4 colocalizes with Rab27 isoforms on SGs and acts as a Rab27 effector (Neeft et al., 2005; Woo et al., 2017), we examined whether Rab27a and Rab27b play distinct roles in vacuole assembly and Munc13-4 membrane retention. Confocal microscopy of cells co-expressing EGFP-Munc13-4 and mCherry-Rab27a confirmed strong colocalization on resting SGs and ionomycin-induced vacuoles, with a Manders’ colocalization coefficient (MCC) of ∼0.65–0.75 (**Fig. 5A, B**). Individual siRNA knockdown of either Rab27a or Rab27b (**Fig. 5C**) completely abolished Ca^2+^-induced vacuole formation assessed by VMAT2 immunofluorescence (**Fig. 5D, E**), indicating that both isoforms are required for vacuole biogenesis.

**Figure 5.**
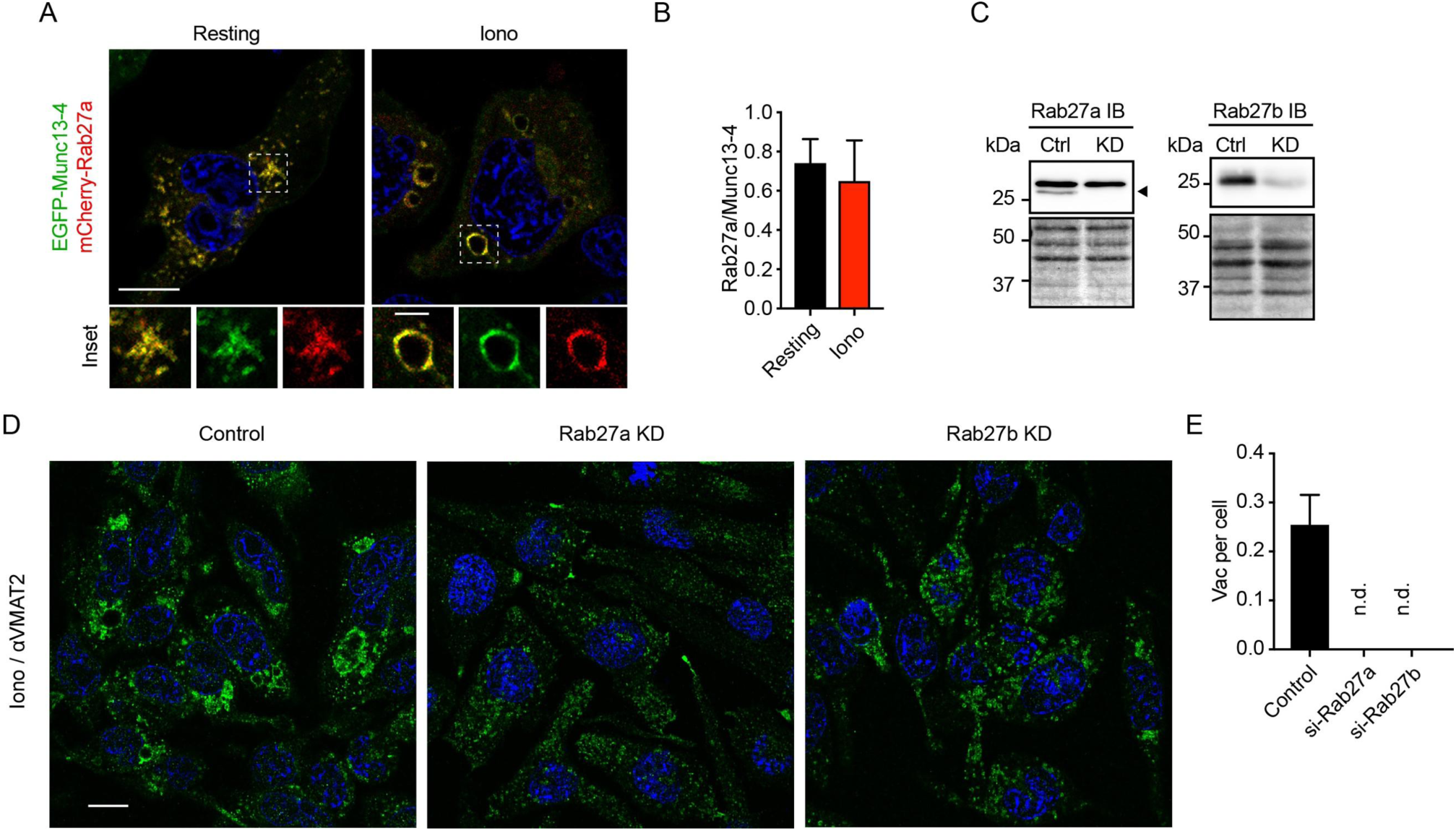
Both Rab27a and Rab27b are required for Ca^2+^-evoked vacuole formation. **A.** Rab27a colocalizes with Munc13-4 on SGs and vacuoles. Representative confocal slices of resting and ionomycin-stimulated RBL-2H3 cells co-expressing EGFP-Munc13-4 (green) and mCherry-Rab27a (red). Scale bars: whole cell, 10 μm; inset, 3 μm. **B.** Manders’ colocalization coefficient (MCC) indicating Rab27a pixels overlapping total Munc13-4 pixels (mean ± SEM, *n* = 6 cells). **C.** Immunoblots verifying siRNA knockdown of Rab27a (arrowhead) and Rab27b relative to control (Ctrl); bottom panels show Ponceau S staining confirming equal protein loading. **D.** Confocal VMAT2 immunofluorescence (green; Hoechst 33342 nuclei in blue) after 20-min ionomycin stimulation in control, Rab27a KD, and Rab27b KD cells. Scale bar, 10 μm. **E.** Vacuole frequency per cell from D (mean ± SEM; *n* = 98, 77, and 78 cells for Control, si-Rab27a, and si-Rab27b, respectively; n.d., not detected).

Strikingly, however, live-cell imaging of resting cells revealed an isoform-specific requirement for membrane retention: siRNA depletion of Rab27a redistributed EGFP-Munc13-4 diffusely into the cytosol, whereas Rab27b knockdown left punctate SG localization intact (**Fig. 6A**). To quantify Munc13-4 membrane exchange dynamics, we performed confocal FRAP (0.28-s intervals) on resting SGs versus ionomycin-induced vacuoles (**Fig. 6B**). Whereas integral membrane CD63-mKate2 showed no recovery after photobleaching, EGFP-Munc13-4 recovered to an identical plateau (∼52% mobile fraction; immobile fraction 0.49 ± 0.10 on SGs versus 0.48 ± 0.07 on vacuoles) within 60 s (**Fig. 6B–D**). Notably, the recovery time constant (τ) was ∼2.5-fold faster on vacuoles (∼7 s) than on resting SGs (∼17 s; *P* < 0.001; **Fig. 6C, D**), demonstrating rapid cytosolic–membrane exchange of Munc13-4 that accelerates on mature vacuoles.

**Figure 6.**
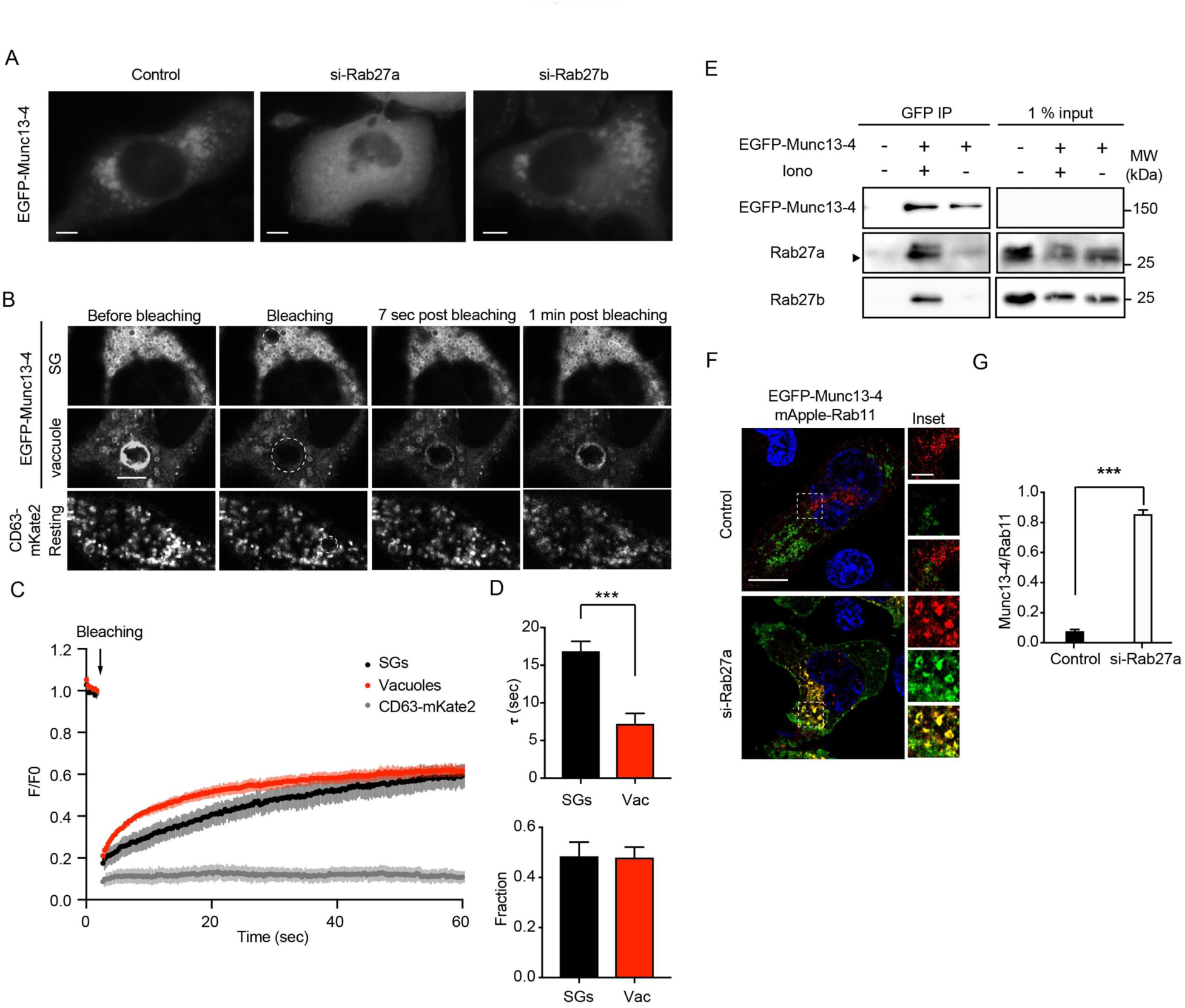
Rab27a mediates basal membrane association of Munc13-4, whereas Rab27a depletion enables switching onto Rab11^+^ vesicles. **A.** si-Rab27a dissociates EGFP-Munc13-4 from SGs into the cytoplasm, whereas si-Rab27b does not. Live-cell snapshots of resting RBL-2H3 cells co-transfected with EGFP-Munc13-4 and control, Rab27a, or Rab27b siRNAs. Scale bars, 5 μm. **B.** Representative confocal FRAP sequences of EGFP-Munc13-4 on resting SGs and ionomycin-induced vacuoles (10 min stimulation), alongside integral membrane CD63-mKate2 on resting SGs. Dashed circles indicate photobleached ROIs. Scale bar, 5 μm. **C.** Normalized fluorescence recovery curves (*F*/*F*_0_) plotted over time (mean ± SEM, *n* = 3 cells per condition). **D.** Recovery time constant τ (s, upper panel; ***, *P* < 0.001) and immobile fraction (lower panel) of EGFP-Munc13-4 on SGs versus vacuoles (mean ± SEM, *n* = 3 cells). **E.** DSP cross-linking GFP-Trap co-immunoprecipitation of endogenous Rab27a (arrowhead) and Rab27b with EGFP-Munc13-4 in resting (−) versus 10-min ionomycin-stimulated (+) cells. **F.** Representative confocal slices showing recruitment of EGFP-Munc13-4 (green) onto mApple-Rab11^+^ vesicles (red) in Rab27a KD cells. Scale bar, 10 μm. **G.** MCC for Munc13-4/Rab11 overlap (superimposed pixels over total Rab11 pixels; mean ± SEM, *n* = 6 cells; ***, *P* < 0.001).

To test whether intracellular Ca^2+^ modulates physical Rab27–Munc13-4 association, cells expressing EGFP-Munc13-4 were cross-linked *in situ* with DSP before GFP-Trap immunoprecipitation. Pull-down of endogenous Rab27a and Rab27b was weak in resting cells but increased markedly following 10-min ionomycin stimulation (**Fig. 6E**). Together with the knockdown data, this indicates that Rab27a maintains basal membrane retention of Munc13-4, while Ca^2+^ elevation reinforces engagement with both Rab27a and Rab27b.

In MDA-MB-231 breast cancer cells, endogenous Munc13-4 is predominantly cytosolic at rest—resembling Rab27a-depleted RBL-2H3 cells—and translocates onto Rab11^+^ endosomes upon Rab11 overexpression or Ca^2+^ elevation (Messenger et al., 2018). Remarkably, expressing mApple-Rab11a in Rab27a-knockdown RBL-2H3 cells redirected cytosolic EGFP-Munc13-4 onto Rab11^+^ vesicles, raising the Munc13-4/Rab11 MCC from <0.10 in control cells to >0.85 (*P* < 0.001; **Fig. 6F, G**). Subsequent ionomycin stimulation further enhanced Munc13-4 recruitment onto Rab11^+^ structures in TIRF planes, as confirmed by overlapping line-intensity scans and frame-subtraction maps (**Supplementary Fig. S4A–C**). Thus, Munc13-4 is not hard-wired to a single Rab partner; rather, relative Rab27a versus Rab11 abundance dictates competitive membrane partitioning of Munc13-4 across cell types.

### Late endosomal SNAREs are required for upstream vacuole assembly

Munc13-4 binds late endosomal (LE) SNARE complexes (Syntaxin-7, Syntaxin-8, Vti1b, and VAMP-8) in RBL-2H3 cells and stimulates Ca^2+^-dependent fusion of LE-SNARE proteoliposomes (Boswell et al., 2012; Woo et al., 2017). Immunofluorescence analysis showed that Syntaxin-7 and VAMP-8 decorated ∼63% and ∼77% of EGFP-Munc13-4^+^ SGs in resting cells, whereas Syntaxin-8 and Vti1b showed <20% overlap (**Fig. 7A, B**). After ionomycin stimulation, all four LE-SNAREs localized to microdomains along the vacuole limiting membrane, with vacuolar coincidence of Syntaxin-8 (∼42%) and Vti1b (∼50%) increasing alongside Syntaxin-7 (∼66%) and VAMP-8 (∼80%) (**Fig. 7C, D**). Individual siRNA depletion (>90% knockdown; **Fig. 7E**) of each LE-SNARE reduced Ca^2+^-evoked Rab27b^+^ vacuole formation by >80% (**Fig. 7F, G**). Because homotypic vacuole biogenesis precedes PM fusion, these data establish a requirement for LE-SNAREs in MVB assembly upstream of exosome release, while leaving open which SNARE/SM complexes directly catalyze terminal MVB–PM pore opening.

**Figure 7.**
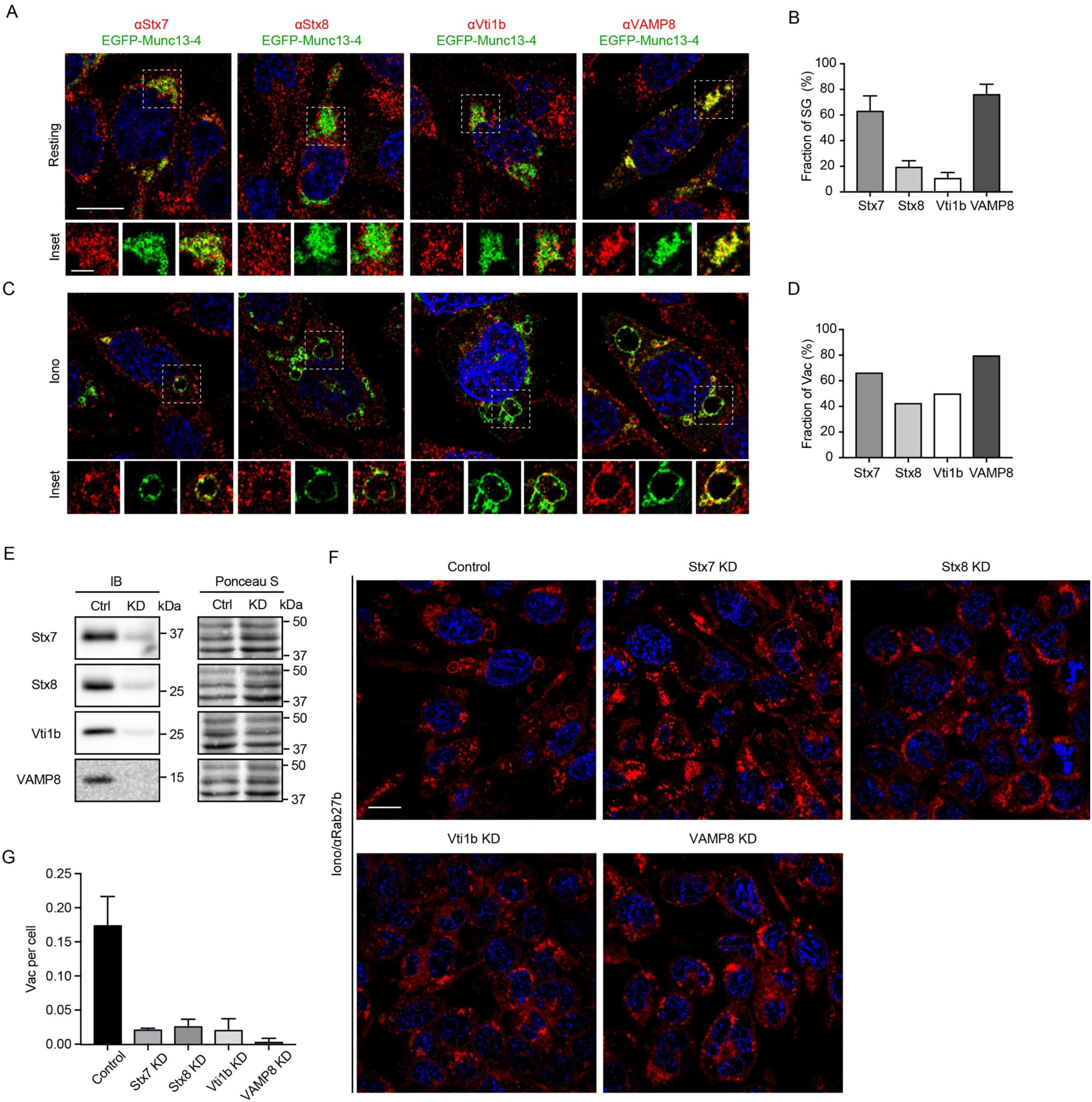
Late endosomal (LE) SNAREs are required for vacuole formation. **A.** Representative confocal slices of resting RBL-2H3 cells expressing EGFP-Munc13-4 (green) immunostained for endogenous Syntaxin-7 (Stx7), Syntaxin-8 (Stx8), Vti1b, or VAMP-8 (red). Scale bars: whole cell, 10 μm; inset, 3 μm. **B.** Percentage of Munc13-4^+^ SGs positive for each LE-SNARE (mean ± SEM, *n* = 3 cells). **C.** Confocal localization of LE-SNARE microdomains on ionomycin-induced vacuoles. **D.** Fraction of vacuoles containing each LE-SNARE on the limiting membrane determined by line-scan analysis (6 confocal slices from 2 cells per SNARE). **E.** Immunoblots verifying individual siRNA knockdown of LE-SNAREs alongside Ponceau S loading controls. **F.** Confocal Rab27b immunofluorescence (red; Hoechst 33342 nuclei in blue) after ionomycin stimulation in control and LE-SNARE KD cells. Scale bar, 10 μm. **G.** Vacuole frequency per cell in control and LE-SNARE KD cells (mean ± SEM; *n* = 154, 179, 295, 299, 232, and 200 cells).

## Discussion

Here, we directly interrogated the pre-fusion maturation and terminal PM fusion stages of Ca^2+^-evoked vacuole exocytosis in live mast cells. Previous live-cell exosome studies relying on pH-sensitive reporters such as CD63-pHluorin have captured terminal luminal neutralization (Messenger et al., 2018; Verweij et al., 2018; Lu et al., 2018) but could not resolve upstream effector retention or stalled post-docking states. Likewise, while Omari et al. (2024) characterized a secretory granule–amphisome exosome route and post-fusion remodeling in RBL-2H3 cells, our simultaneous epi-TIRF and staged inhibitor experiments pinpoint the terminal Munc13-4-dependent transition into fusion pore opening and stabilization. Crucially, our findings indicate that terminal MVB exocytosis proceeds as a dynamic continuum rather than segregating into rigid biochemical boxes of ‘tethering’ versus ‘priming’: when Munc13-4 is acutely inhibited after vacuole assembly, vacuoles still contact the basal PM footprint, yet fusion pore opening fails and nascent pores prematurely collapse (**Fig. 4**). Without overturning prior roles of Munc13-4 in granule tethering (Elstak et al., 2011; Chicka et al., 2016), these observations demonstrate that terminal pore opening and dilation are the steps most acutely sensitive to Munc13-4 perturbation.

### Vacuoles as a model system for MVB maturation, cargo convergence, and fusion pore dynamics

Enlarged vacuoles in RBL-2H3 cells permitted direct optical measurement of MVB–PM fusion pore diameters (1.4 ± 0.5 μm), open-pore lifetimes (2.38 ± 1.63 min), exocytic organelle shrinkage without full membrane collapse, and CD63^+^ exosome expulsion confirmed by HPF-EM (**Fig. 1**). Moreover, vacuoles served as a convergent sorting reservoir integrating distinct cargo routes: endogranular CD63 coalesced via homotypic SG/endosome fusion, whereas PM-resident CD9 was internalized into small endocytic vesicles following initial SG exocytosis before merging into maturing vacuoles to bud inward as ILVs (**Fig. 2F–H**; Pols and Klumperman, 2009; van Niel et al., 2011; Bobrie et al., 2012; Kowal et al., 2016). Although RBL-2H3 cells are a tumor-derived mast cell line—warranting future corroboration in primary mast cells—antigen-stimulated primary mast cells exhibit analogous compound exocytosis and MVB enlargement, and Ca^2+^-regulated convergence of Rab7^+^ and Rab11^+^ compartments is conserved across cytotoxic T cells, erythroleukemia cells, neurons, and cancer cells (Raposo et al., 1997; Savina et al., 2003, 2005; Ménager et al., 2007; Lachenal et al., 2011; Woo et al., 2017; Messenger et al., 2018).

### Abundance-weighted Rab partitioning reconciles biochemical and cellular Ca²⁺ sensitivity

Munc13-4 interacts with multiple Rab isoforms—including Rab11, Rab15, Rab27a, Rab27b, and Rab37— across diverse secretory lineages (Shirakawa et al., 2004; Neeft et al., 2005; Ménager et al., 2007; Pivot-Pajot et al., 2008; Wood et al., 2009; Zografou et al., 2012; Singh et al., 2013; Higashio et al., 2016; Johnson et al., 2016). Our FRAP, DSP cross-linking, and Rab-switching experiments (**Figs 5, 6**; **Supplementary Fig. S4**) reveal that Munc13-4 undergoes rapid cytosolic–membrane exchange (τ ≈ 7 s on vacuoles) and partitions competitively between Rab27a and Rab11 according to their relative cellular abundance. High endogenous Rab27a retains Munc13-4 on the SG/vacuole pathway in mast cells, whereas depleting Rab27a allows Rab11 to capture Munc13-4 onto Rab11^+^ compartments in a Ca^2+^-enhanced manner indistinguishable from MDA-MB-231 breast cancer cells (Messenger et al., 2018).

Notably, this rapid-exchange, abundance-weighted Rab partitioning model also resolves a longstanding quantitative discrepancy between biochemical reconstitution and intact-cell physiology. In reconstituted liposome assays, Munc13-4 engages PS/PE-containing membranes with steep, high-affinity Ca^2+^ sensitivity (apparent *K*_d_ < 50 μM; Boswell et al., 2012). By contrast, Ca^2+^-evoked secretion in intact RBL-2H3 cells exhibits a markedly shallower dependence (EC_50_ ≈ 0.5 mM, Hill coefficient ≈ 1.3; Woo et al., 2017). Our data indicate that intrinsic Ca^2+^–lipid binding capacity is necessary but not sufficient in cells; because the pool of Munc13-4 available at fusion-competent membranes is constrained by competitive, rapid-exchange partitioning across Rab27a, Rab27b, and Rab11 compartments, Rab competition tempers steep biochemical lipid binding into a graded physiological secretory response.

### Downstream SNAREs and therapeutic implications of context-specific Rab–Munc13-4 coupling

While canonical LE-SNAREs (Syntaxin-7, Syntaxin-8, Vti1b, and VAMP-8; Prekeris et al., 1999; Antonin et al., 2000) proved essential for upstream Ca^2+^-triggered vacuole assembly (**Fig. 7**), defining the exact *trans*-SNARE/SM complex (potentially involving Syntaxin-11/Munc18-2, SNAP-23, or tethering complexes such as HOPS) acting at the final MVB–PM pore will require acute peptide/toxin perturbation or high-resolution structural analysis.

More broadly, distinguishing universal from cell-type-specific exosome regulators carries direct therapeutic significance. Metastatic cancer cells upregulate Munc13-4 via TGF-β signaling to drive Rab11-coupled secretion of pro-metastatic exosomes (Messenger et al., 2018), while its C2-domain paralog BAIAP3 regulates related secretory and exosomal pathways (Zhang et al., 2017; Lu et al., 2018). Importantly, recent discovery of small-molecule inhibitors that selectively disrupt the Munc13-4–Syntaxin-7 interface to suppress endosomal TLR-driven systemic inflammation without broad cellular toxicity provides proof of concept that individual Munc13-4 partner interfaces are pharmacologically separable (Johnson et al., 2026). Because innate immune cells rely primarily on the Rab27a–Munc13-4 interface whereas cancer cells exploit the Rab11–Munc13-4 axis, selectively targeting context-dependent Munc13-4 conformations or specific Rab–Munc13-4 interfaces offers a rational path to suppress pathological tumor exosome release while sparing host immune defense.

## Materials and Methods

### Cell culture, transfection, and lentiviral transduction

RBL-2H3 cells (CRL-2256; ATCC, Manassas, VA) were cultured in Eagle’s Minimum Essential Medium (MEM) supplemented with 15% fetal bovine serum (FBS) and Antibiotic-Antimycotic (Life Technologies, Carlsbad, CA) at 37°C under 5% CO_2_. For live-cell and immunofluorescence microscopy, glass coverslips were coated with 0.1 mg/ml poly-D-lysine and 30 μg/ml bovine fibronectin (Sigma-Aldrich, St. Louis, MO) at 37°C. For electroporation, 2 μg plasmid DNA and/or 2 μM siRNA duplexes (final concentration) were mixed with RBL-2H3 cells in Cytomix buffer supplemented with 5 mM glutathione and 2 mM ATP (van den Hoff et al., 1992) and pulsed twice (1 ms, 225 V) in an ECM 830 electroporator (Harvard Apparatus, Holliston, MA). Culture medium was replaced after 24 h and assays were conducted 48 h post-transfection. Stable Munc13-4 knockdown cells were generated by lentiviral shRNA transduction as previously described (Woo et al., 2017; Messenger et al., 2018) and verified by anti-Munc13-4 and anti-GAPDH immunoblotting.

### Antibodies, plasmids, and chemical reagents

Goat and rabbit polyclonal anti-Munc13-4 antibodies and anti-VMAT2 immunofluorescence reagents were used as previously described (Woo et al., 2017). Rabbit polyclonal anti-Rab27b (168103) was from Synaptic Systems (Göttingen, Germany). Sheep polyclonal anti-Rab27a (AF7245) and antibodies against Syntaxin-7 (AF5478), Syntaxin-8 (AF5448), and VAMP-8 (AF5354) were from R&D Systems (Minneapolis, MN). Rabbit monoclonal anti-CD9 (EPR2949) was from Abcam (Cambridge, UK); mouse monoclonal anti-CD63 (H5C6) and anti-GM130 were from BD Biosciences (San Jose, CA); anti-Vti1b (1410097) and anti-GAPDH were from Sigma-Aldrich. Alexa Fluor-conjugated secondary antibodies were from Molecular Probes/Thermo Fisher Scientific (Eugene, OR). Bexin-1 and its inactive analog Bexin-5 were characterized previously (Bruinsma et al., 2018). mApple-Rab11a was a gift from Michael Davidson (Addgene plasmid #54942); EGFP-Munc13-4, mCherry-Rab27a, EGFP-Rab7, CD63-mKate2, CD63-tGFP, CD9-mCherry, and CD9-EGFP were generated as described previously (Woo et al., 2017; Messenger et al., 2018).

### Immunofluorescence microscopy

Cells were washed with PBS, fixed in 4% formaldehyde in PBS for 30 min at room temperature, and permeabilized in 0.2% Triton X-100 in PBS for 20 min at room temperature. After blocking for 1 h in 5% FBS, goat serum, or donkey serum, coverslips were incubated overnight at 4°C with primary antibodies in blocking buffer, washed, and incubated for 1 h at room temperature with 20 μg/ml Alexa Fluor-conjugated secondary antibodies. Nuclei were stained with 2 μg/ml Hoechst 33342 for 5 min, and coverslips were mounted in SlowFade Gold antifade reagent (Life Technologies).

### Simultaneous epi-TIRF microscopy, FRAP, HPF-EM, and image quantification

Live-cell imaging was performed at 37°C in a humidified stage-top chamber (Tokai Hit, Shizuoka, Japan) in Tyrode’s buffer containing 10 mM CaCl_2_ with 2 or 2.5 μM ionomycin as indicated. Simultaneous epifluorescence and TIRF time-lapse series were acquired at 20-s intervals (or 10-s intervals for rapid cargo tracking in Fig. 2G) on a Nikon Eclipse Ti microscope equipped with an iXon Ultra EM-CCD camera (Andor), a Nikon APO 100× TIRF objective (NA 1.49), and a Perfect Focus System (PFS) switching focal planes between the cell body and basal PM in <1 s. High-pressure freezing, freeze substitution, and transmission electron microscopy were conducted as described previously (Woo et al., 2017). Epifluorescence images were deconvolved in NIS-Elements (Nikon). Colocalization (Manders’ colocalization coefficient, MCC) was analyzed using the GDSC plugin (University of Sussex, UK) for confocal stacks and JACoP in ImageJ/Fiji for deconvolved images. Radial intensity profiles across vacuoles and fusion pores were measured using concentric 1-pixel radial sweeps in ImageJ (Radial Profile Angle plugin; Woo et al., 2017). For FRAP analysis, circular ROIs on resting SGs or ionomycin-induced vacuoles (10 min stimulation) were photobleached with 100% 488-nm laser power on a Nikon A1 confocal system and imaged continuously at 0.28-s intervals; recovery time constants (τ) and immobile fractions were fitted in NIS-Elements (Reits and Neefjes, 2001).

### Exosome secretion assays and sucrose density gradient purification

For 96-well fluorometric secretion assays, RBL-2H3 cells expressing CD63-mKate2, CD9-EGFP/GFP, or soluble GFP control were plated in triplicate and stimulated 48 h later with 2 μM ionomycin in Tyrode’s buffer at 37°C for 30 min (or indicated time points). Supernatants and 1% Triton X-100 cell lysates were measured to calculate % secretion relative to total cellular fluorescence (Woo et al., 2017). For endogenous exosome slot-blot assays (Messenger et al., 2018), cells at ∼70% confluency were washed with PBS containing 100 μM EDTA and stimulated in phenol red-free DMEM containing 10 mM CaCl_2_ and 2.5 μM ionomycin for 20 min at 37°C (with 10 μM Bexin-1 or Bexin-5 added 10 min prior where indicated). Conditioned media were cleared sequentially at 300 × *g* (5 min), 1,000 × *g* (10 min), and 10,000 × *g* (10 min), vacuum-filtered onto nitrocellulose membranes via a slot-blot apparatus, and immunoblotted for CD63, CD9, and *cis*-Golgi marker GM130 alongside 1% total cell lysates.

For sucrose density block fractionation (Lötvall et al., 2014; Shurtleff et al., 2016), conditioned media from large-scale cultures stimulated with or without 2.5 μM ionomycin were cleared at 300 × *g* (5 min), 1,000 × *g* (10 min), and twice at 10,000 × *g* (10 min), pelleted twice at 100,000 × *g* for 120 min, and layered onto a stepwise sucrose gradient (0%, 20%, 40%, and 60% sucrose). Vesicles banding at the 20%/40% sucrose interface were harvested, lysed in 4× SDS sample buffer (non-reducing without DTT for CD63; reducing with DTT for other markers), and analyzed by immunoblotting.

### In situ cross-linking immunoprecipitation

RBL-2H3 cells expressing EGFP-Munc13-4 were rested or stimulated with ionomycin in Tyrode’s buffer for 10 min at 37°C, washed with PBS, and cross-linked with dithiobis(succinimidyl propionate) (DSP; Thermo Fisher Scientific) for 30 min at room temperature. After quenching/washing in PBS, cells were solubilized in ice-cold 1% Triton X-100 in PBS, clarified, and incubated with anti-GFP nanobody magnetic beads (GFP-Trap; ChromoTek, Martinsried, Germany). Washed immune complexes were eluted and resolved by SDS-PAGE for immunoblotting against EGFP-Munc13-4, Rab27a, and Rab27b (Woo et al., 2017).

### Statistical analysis

Data are presented as mean ± SEM, mean ± SD, or median with interquartile range (IQR) as specified in each figure legend. Statistical comparisons were performed using unpaired two-tailed Student’s *t*-tests or two-sided Fisher’s exact tests (for categorical pore-collapse frequencies) in GraphPad Prism or Microsoft Excel (*, *P* < 0.05; **, *P* < 0.01; ***, *P* < 0.001).

## Acknowledgements and Declarations

### Acknowledgements

Experimental data were generated by S.S.W. in the Department of Biochemistry at the University of Wisconsin–Madison.

### Author Contributions

S.S.W. conceived the study, designed and performed all experiments, analyzed the data, and wrote the manuscript.

---

**Figure S1. Related to Figs 1 and 2.**
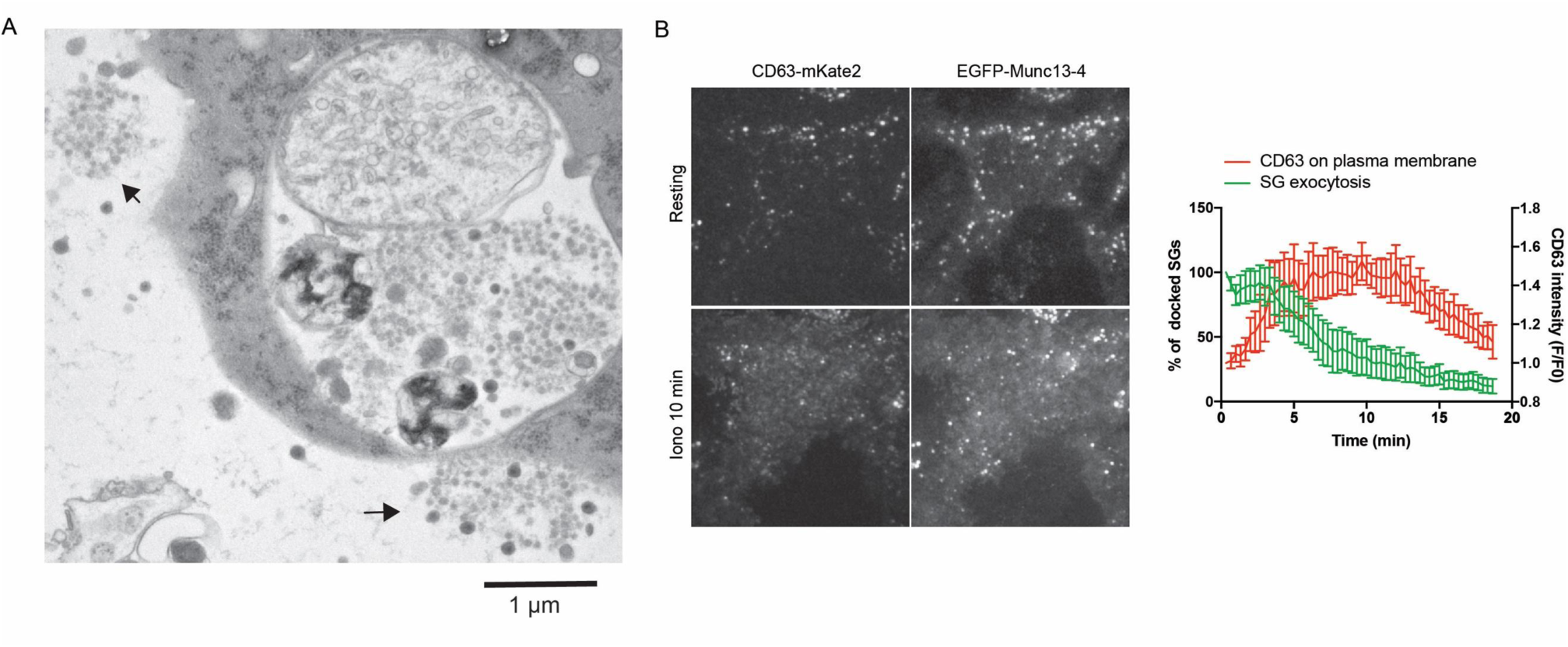
**A.** HPF-EM micrograph showing aggregated exosomes (arrows) released from ionomycin-stimulated RBL-2H3 cells. Scale bar, 1 μm. **B.** Kinetics of SG exocytosis and CD63 insertion into the plasma membrane. Dual-channel TIRF images (left) and quantification (right; mean ± SEM, *n* = 7 cells) showing loss of docked SGs in the TIRF field (green, % of initial SG number) and a concurrent ∼40% increase in basal PM CD63-mKate2 fluorescence intensity (red, *F*/*F*_0_) within 5 min after ionomycin stimulation.

**Figure S2. Related to Fig. 2.**
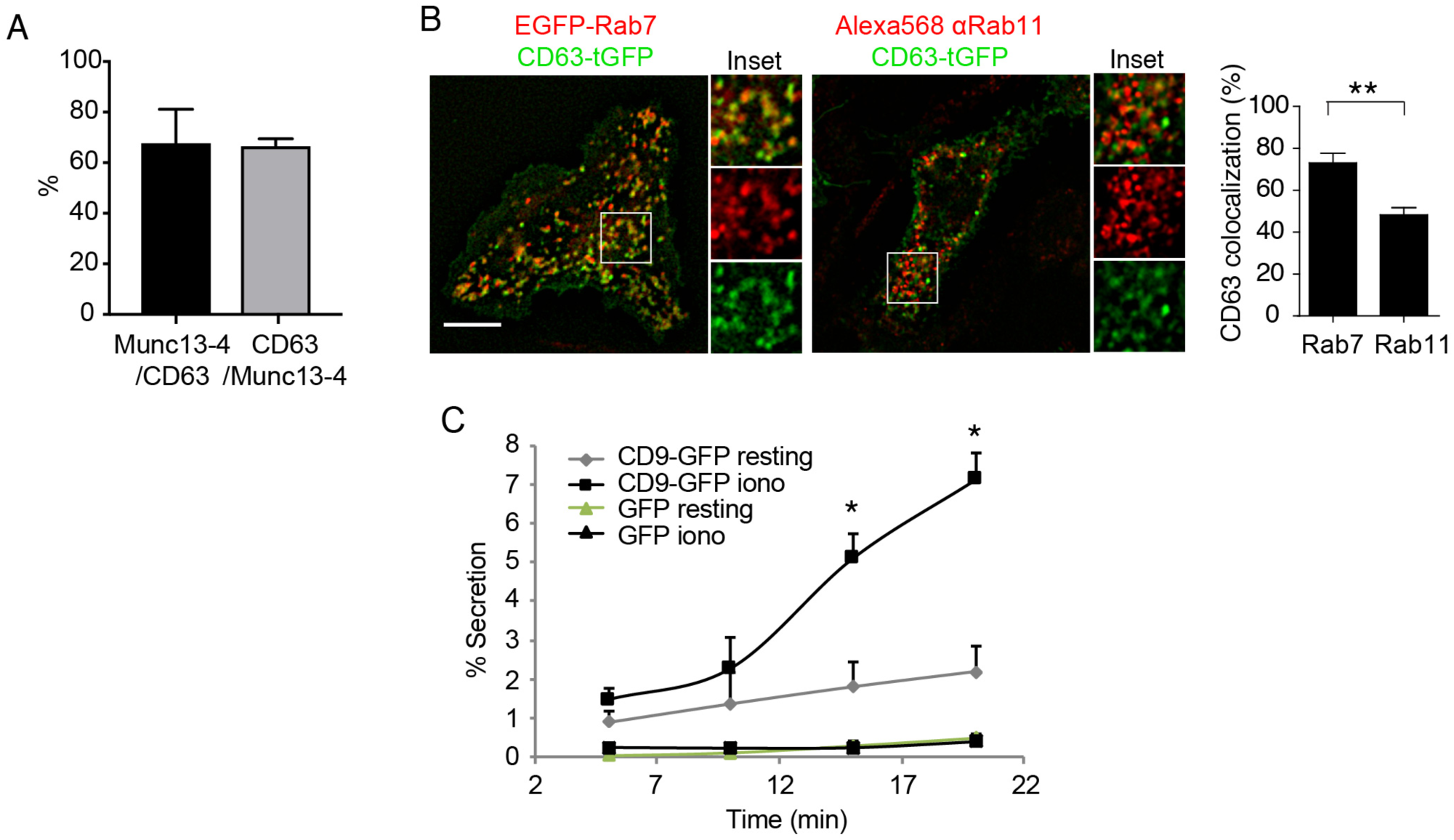
**A.** Line-scan colocalization (%) between ring-shaped EGFP-Munc13-4 and CD63-mKate2 structures (*n* = 6 cells from 2 independent experiments). **B.** Epifluorescence images (left; scale bar, 10 μm) and quantification (right; mean ± SEM, *n* = 6 cells; **, *P* < 0.01) of CD63-tGFP (green) colocalization with EGFP-Rab7 or anti-Rab11 immunofluorescence (red). **C.** Time course of ionomycin-stimulated secretion (% of total) in cells expressing CD9-GFP versus soluble GFP negative control (mean ± SEM, *n* = 5; *, *P* < 0.05 comparing CD9-GFP iono versus GFP iono).

**Figure S3. Related to Fig. 3.**
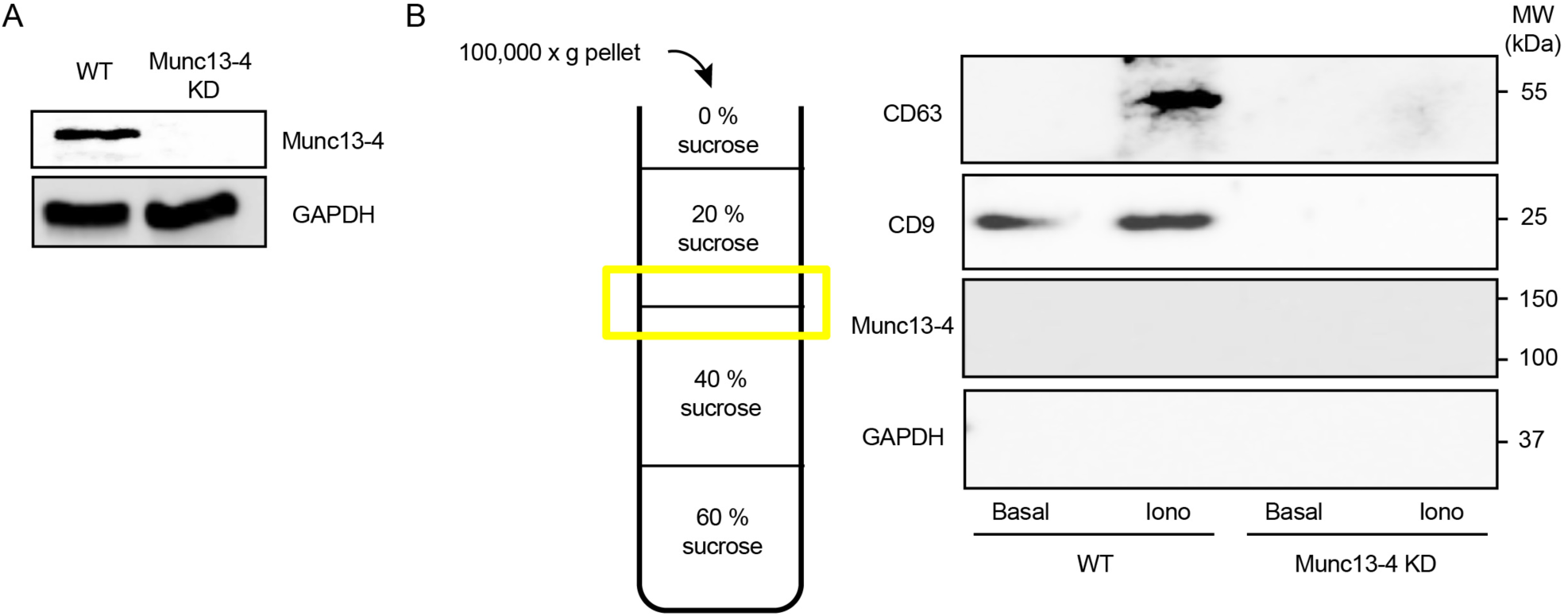
**A.** Immunoblot confirming lentiviral shRNA knockdown of Munc13-4 in RBL-2H3 cells with GAPDH as a loading control. **B.** Stepwise sucrose density block fractionation of exosomal CD63 and CD9 from WT and Munc13-4 KD cells stimulated with or without 2.5 μM ionomycin for 30 min. Left: Schematic showing collection of the 20%/40% sucrose interface from 100,000 × *g* pellets. Right: Immunoblots for CD63, CD9, Munc13-4, and GAPDH (Messenger et al., 2018).

**Figure S4. Related to Fig. 6.**
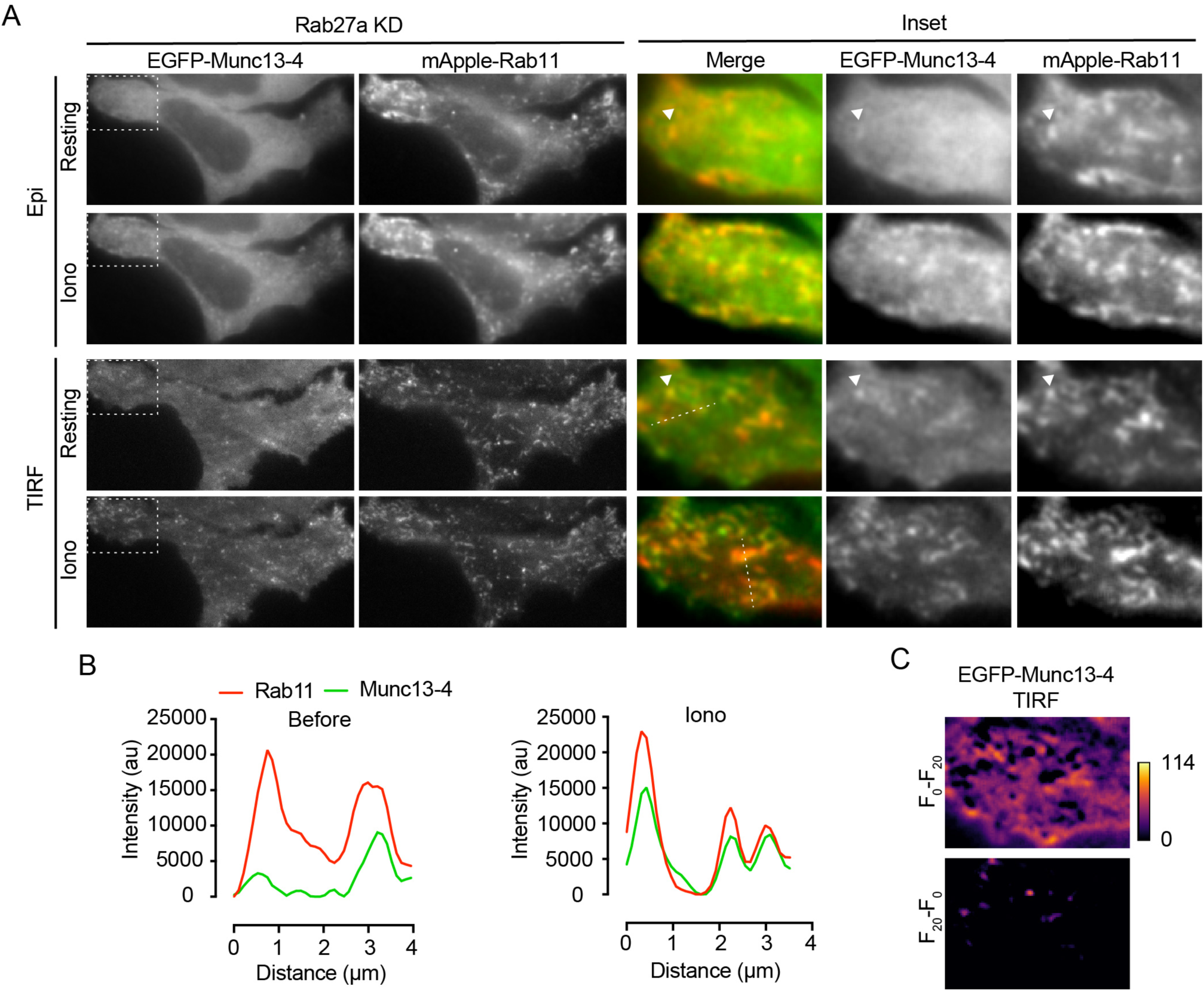
Munc13-4 in Rab27a KD cells translocates to Rab11^+^ vesicles in a Ca^2+^-dependent manner. **A.** Simultaneous Epi and TIRF images of Rab27a KD cells co-expressing EGFP-Munc13-4 and mApple-Rab11 before (Resting) and after ionomycin stimulation (Iono). Arrowheads mark Rab11^+^ vesicles associated with Munc13-4. **B.** Line-scan intensity profiles along the dashed TIRF lines in A showing Ca^2+^-enhanced overlap of EGFP-Munc13-4 with Rab11^+^ puncta. **C.** Frame-difference heatmaps (*F*_0_ − *F*_20_, upper; *F*_20_ − *F*_0_, lower) of TIRF EGFP-Munc13-4 intensity demonstrating rapid cytosolic depletion and vesicular recruitment within 20 s of stimulation.

